# Whole brain evaluation of cortical micro-connectomes

**DOI:** 10.1101/2022.10.05.510240

**Authors:** Kouki Matsuda, Arata Shirakami, Ryota Nakajima, Tatsuya Akutsu, Masanori Shimono

## Abstract

The brain is an organ that functions as a network of many elements connected in a non-uniform manner. Especially, the cortex is evolutionarily newest, and is thought to be primarily responsible for the high intelligence of mammals. In the mature mammalian brain, all cortical regions are expected to have some degree of homology, but have some variations of local circuits to achieve specific functions enrolled by individual regions. However, few cellular-level studies have examined how the networks within different cortical regions differ. This study aimed to find rules for systematic changes of connectivity (microconnectomes) across 16 different cortical region groups. We also observed unknown trends in basic parameters *in vitro* such as firing rate and layer thickness across brain regions. The results revealed that the frontal group shows unique characteristics such as dense active neurons, thick cortex and strong connections with deeper layers. This suggests the frontal side of the cortex is inherently capable of driving, even in isolation.

This may suggest that deep layers of frontal node provide the driving force generating a global pattern of spontaneous synchronous activity, such as the Default Mode Network. This finding may explain why disruption in this region causes a large impact on mental health.

## Introduction

The brain is an organ with very nonuniformly connected components working together as a network. However, the components also have unique characteristics as individual brain regions. If we are able to extract any systematic rule about such nonuniformity in terms of internal characteristics of local components, it may provide important insights about how the brain processes information in parallel across different scales.

### Focus on Cortex

Within the brain, the neocortex is an evolutionary novelty and an important component of the brain that supports the high intelligence of mammals, including humans [Kaas, 2020]. At first glance, it is difficult to distinguish unique characteristics of each subregion because the neocortex has a simple structure in which similar circuits are arranged throughout the entire cortical sheet.

However, although there is some degree of similarity, there is also a certain degree of uniqueness [DeFelipe, 2002; Nelson, 2002; Douglas, Martin, 2004]. In fact, in the brain of a mature animal, each cortical region is assigned a distinct set of functions. To perform different functions, individual brain regions must either receive different inputs from the outside via the macroconnectome or/and have different local circuits (microconnectomes) within them.

### Macro- Micro- and Meso-scales

There has recently been extensive progress in the analysis of structural and functional connectivity topologies between macroscopic cortical regions [Sporns, 2011]. In the case of human subjects, the Human Connectome Project has produced a number of important results [van Essen et al., 2013; Toga et al., 2012]. For example, it is already known that brain regions are non-uniformly connected to each other, and that their patterns of connectivity are also closely related to their functions [Bullmore, Sporns, 2012 ; Gu et al., 2015; Fornite, et al. 2016; Deco et al., 2011]. In non-human animals, the macro-connectome is being studied with increasing precision [Oh et al., 2014; Hintiryan, 2016; Stafford et al., 2014; Goulas et al., 2019; Hayashi et al., 2021]. Information about the network structure is also used for simulation of whole brain dynamics, and various platforms of such simulation are being created [Ritter et al., 2013]. It is also expected to be applied to the comparison of healthy and diseased groups [Aerts et al., 2018; Jirsa et al., 2017; Melozzi, et al., 2019].

In addition to these macro scales, the micro scale is also important. If one wants to accurately reproduce some real quantities via a computational model, we need to carefully consider local circuits. This is because while the broad content of visual, auditory or motor behavior is represented by individual global brain regions, the detailed content, such as fine and smooth movements, is highly dependent on local circuits [Wessberg et al., 2000; Camena et al., 2003; Kiani, et al., 2007; Nicolelis, Lebedev, 2009].

Moreover, the microscale and macroscale features of the cortex cannot be separated. The connections between macroscopic cortical regions are closely related to the laminar structure of individual cortical regions. This is because the hierarchy between cortical regions is defined by the differences in the layer structure of each region [Felleman, Van Essen, 1991]. Furthermore, the mesoscale network existing between the two scales is thought to be less understood than other scales [Buzaki, Christen, 2016; Kennedy et al.] One of the reasons why this scale is less understood because the network of circuits at that scale is the most complex. This also suggests that this scale may encodes much information. In an effort to bridge these scales, some recent simulation studies have begun to include not only neuron density in individual regions but also microscopic connections within individual brain regions [Markram et al., 2015; Schiner et al., 2021].

Although experimental data has begun to provide information on the neuron density and cortical layer thickness of each brain region, it has not yet been possible to understand how the topology of cellular connections within each brain region differs between brain regions and to use this data to inform simulations of the entire brain. In fact, a major challenge is that such data is not sufficiently available over a wide areas of the cortex.

### Inter-regional similarities and differences of cortical micro-circuits

In the past, detailed comparisons of the structure of local circuits in different regions have been mainly made among a few cortical areas [Nelson, 2002]. It has been suggested that cortical local circuits share a common design and constitute a “canonical circuit” [Miller, 2016]. At the same time, each region also has a distinct profile of connectivity [De Biase, L. M., Bonci, A. 2019; Batiuk et al., 2020; Harris, Shepherd, 2015]. In other words, each area can have systematic differences according in the spatial distances and topological distances of their connections [DeFelipe, 2002; Wang et al., 2020].

Collaborative evidence has come from comparisons of cell density and cortical thickness in several regions. For example, Fulcher *et al*. reported topographical similarities in large gradients of cell structure, gene expression, density of interneurons, and long-range axonal connectivity from primary sensory cortex to prefrontal cortex in the mouse cerebral cortex [Fulcher et al., 2019]. Studies in the broader mammalian species have also shown that cell density varies across cortical regions, a finding that is being rapidly elucidated not only by classical immunostaining [Collins et al., 2010; Herculano-Houzel et al., 2013; Keller et al., 2018], but also by emerging transparency techniques [Hama et al., 2011; Susaki et al., 2014; Chung et al., 2013]. Systematic rules for the relationship between connections between brain regions and cell density are also becoming clearer [Herculano-Houzel et al., 2013; Shimono, 2013].

In spite of such a wide interest and a huge amount of research, there are still few global and quantitative evaluations of how >100 neurons internally exchange information among different categories in local neural circuits within each brain region based on experimental data. In particular, there are almost no studies that have made a wide comparison between brain regions, stepping into the topological differences in functional connectivity. Therefore, if we can extract the rules of “gradual systematic differences of individuality”, such information is expected to be a source of significant benefits not only for brain modeling but also for, in a wide range of aspects such as contrast with other non-uniformity indicators, understanding of disease states, etc.

Approaches to exploring the topology of connections between hundreds or more neurons have evolved into a research field called the microconnectome [Schröter et al., 2017].

However, all previous work on the microconnectome and related studies have been confined to a limited number of brain regions and have not made comparisons between brain regions [Lefort, et al., 2009; Lubke, Feldmeyer, 2007; Ko et al., 2013]. Our past studies have also focused on a single brain region to study the micro connectome, and have repeatedly evaluated and improved the technology [Kajiwara et al., 2021; Shimono, Beggs, 2014].

This present study extended the technique to all cortical regions of the mouse, and evaluated how each cortical brain region differs especially in terms of the topological differences of functional local circuits. Prior to this, we also evaluated some basic parameters such as firing rate and layer thickness, which are not yet fully known from previous studies. Here we evaluated density and functional interconnections from active neurons, something that cannot be captured by applying staining technique to dead brain slices.

In our results, we noticed a significant trend in the strength of functional connections within the deep layers of brain regions that were categorized into frontal groups. We can expect the strength of the connections in a region quantitatively reflect which of the internal connections are prioritized information flows. For example, the information that flows in through the external structural connections may represent the default mode network.

## Results

We aimed to compare the differences in functional networks between neurons in brain regions. Before that, we also compared basic physiological or anatomical metrics such as age, cortical thickness and neuron densities. Therefore, we divided the brain into 16 circumferential categories based on the rotating angle as shown in fig.2-a, and defined abbreviated names as listed in table 1. In the following manuscript, we simply call the 16 individual brain region categories as given names based on the angle.

**Table 1:** Definition of 16 regional groups

|  | L(Left) | R(Right) |
| --- | --- | --- |
| OD | L Occipital Dorsal | R Occipital Dorsal |
| D | L Dorsal | R Dorsal |
| FD | L Frontal Dorsal | R Frontal Dorsal |
| F | L Frontal | R Frontal |
| FV | L Frontal Ventral | R Frontal Ventral |
| V | L Ventral | R Ventral |
| OV | L Occipital Ventral | R Occipital Ventral |
| O | L Occipital | R Occipital |

### Basic metric 1: Age

Since the neuron density, firing rate, and network architecture are likely to change at different ages, it is necessary to measure all data for the same age group to simply compare between brain regions.

First of all, we show the age of individuals included in the 16 circumferential categories (fig. 2-b). The average age of each category shows that they are almost stable at around 4 weeks old (3∼5 weeks).

**Figure 1.**
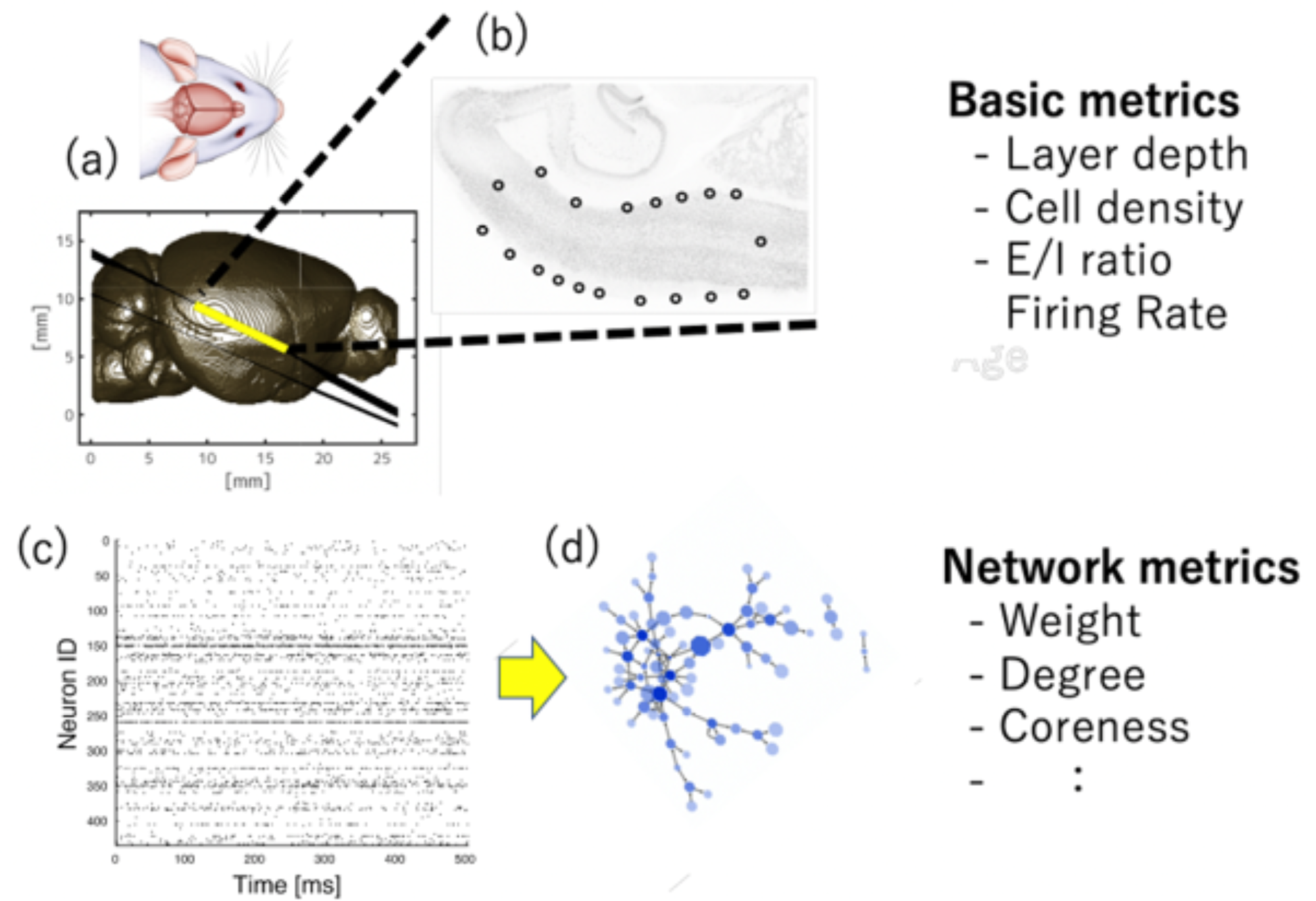
Scheme of the whole analysis: (a) An example of a mouse brain magnetic resonance (MR) images. We cut out acute slices at the location of the lines, and measured electrical activity from them, and then immunostained them. (b) An example of immunostaining. The cortical area surrounded by black circles indicates the area where electrophysiological measurements were also performed. (c) Electrophysiological measurements provide a time series of spikes in the activity of hundreds of neurons. (d) From the time series, causal interactions were inferred and effective networks were constructed. We evaluated not only basic physiological metrics, but also various metrics of the effective networks.

**Figure 2.**
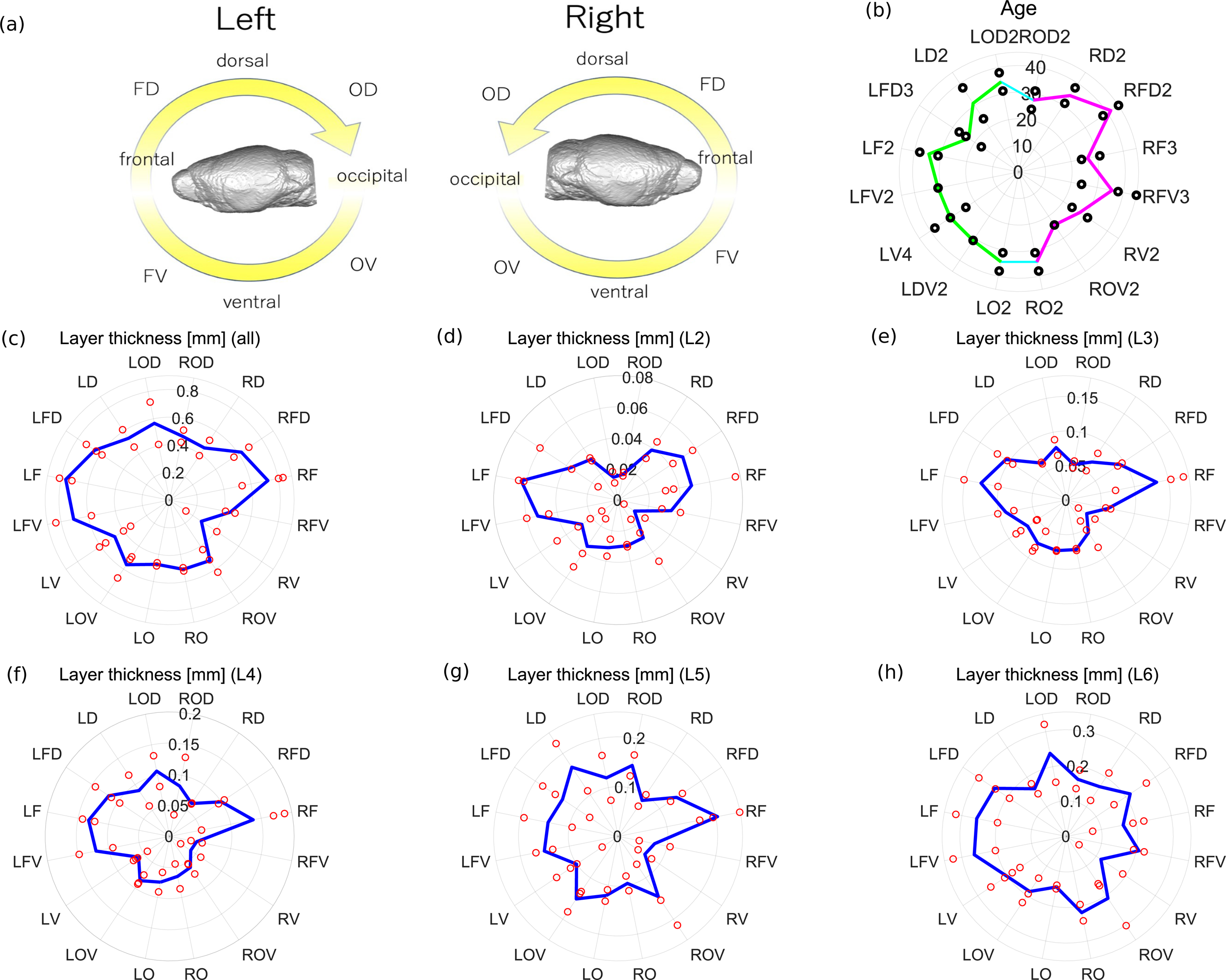
Radar charts of age and cortical thickness: (a) This panel expresses classification of the brain in 16 categories and their abbreviation names. Table 1 refers to these definitions of abbreviation names. In the following seven panels, the left hemisphere corresponds to the left half of the circle, and the right hemisphere to the right half. (b) This panel summarizes ages of mice used in individual experiments. Lines show the averages of the data within each of the 16 categories. (The left and right hemispheres are expressed with red and blue colored lines, respectively.)Black points show the age of each individual mouse used in this experiment. (c) represents the thickness of all layers of the cortex, the line is the average value for each angle, and black dots express raw data. In addition to the panel (c), panels (d)∼(h) express the thickness of individual layers (layer 2, 3, 4, 5, 6) separately.

### Basic metric 2: Layer thickness

Next, we observed some basic anatomical or physiological indices according to two steps. First, we compared the cortical thickness among the 8 categories for individual hemispheres (fig. 2-b). Previous studies reported that cortical regions categorized as frontal group (or group F in our definition) are thicker than other regions [Pagani, et al. 2016; garand’Malson et al. 2013].

Therefore, we performed the following statistical tests, hypothesizing that 2-3 adjacent brain regions around the frontal group would also show differences compared to other regions. Here, we found that the cortical thickness was higher in an adjacent pair of brain regions, categorized into the frontal and frontal dorsal groups, than other remaining 7 groups (p<0.005, Mann-Whitney test, groups F and FD) commonly between two hemispheres.

Second, we extended this finding further by observing the thickness of individual layers (fig. 2-c). Here, we found layer 4 was observed to be thinnest on the dorsal side, which corresponds to the motor cortex. This finding was also consistent with arguments from previous studies that stated layer 4 is not prominent in many regions, for example, the motor cortex [Yamakawa et al. 2014; Garcia-Cabezas et al. 2014].

More interestingly, this trend of selectively increasing thickness in the F and FD groups is common across the many layers (Layer2∼5, p<0.05, Mann-Whitney test), except in the deepest layer (Layer 6, p>0.05, Mann-Whitney test). Here, the statistical test was performed based on the same hypothesis for the case of all layers. We will discuss this new finding in the discussion section.

### Basic metric 3: Density of active neurons

Next, we observed the neuron density. Density of active neurons is calculated by dividing the number of neurons with a firing rate greater than 0.1Hz by the area of the region where the neurons were present.

Here, notice that we produced acute slices that were coronal, and not tangential, to the brain surface. Therefore, the thickness of the cortex could correspond with one axis of our electrode array. Besides, we defined E/I (excitatory or inhibitory) categories for individual neurons based on properties of output connections of the neurons [Kajiwara et al. 2021]. Therefore, we could calculate the density of active excitatory and inhibitory neurons by dividing the numbers of identified excitatory or inhibitory neurons by the spatial size covering a region of focus. Here, we put slices on a rectangular recording region to keep the cortical surface closely in parallel with one axis of the recording region. Therefore, for simplicity, we were also able to qualitatively use cortical thickness as the normalization factor instead of the size of the recording cortical region.

Here, we hypothesized again that a few groups near the frontal region should be the main target of the statistical test to compare with other groups. Then, for the density of active excitatory neurons for all layers, frontal and frontal ventral regions showed significantly higher values than other regions (p<0.05 (p=0.0419), Mann-Whitney test).

Additionally, we performed a separate analysis of the individual layers defined based on immunostaining. The results showed that the trend of non-uniformity of density of active excitatory neurons was completely different between superficial and deep layers.

Specifically, in deep layer 5 or 6, the high density around the frontal side was pronounced (Layer2∼4, p>0.05), while, in (deeper?) layers, the high density around the frontal side was not so strongly observed (Layer5, p<0.05, p=0.035; Layer6, p<0.05, p=0.015, F & FV).

Briefly speaking, the trend, that the density of active excitatory neurons is high around the frontal side observed for all layers, is influenced by the same but much stronger trend observed in the deep layers.

### Basic metric 4: E/I balance

We aimed to explore the differences between cortical layers in addition to obtaining a distinction between excitatory and inhibitory neurons.

Past studies have shown that E/I balance is well mixed and distributed in different areas of the cortex [Xue et al., 2014; Moreau et al., 2010]. We quantified E/I balance as the ratio between numbers of active excitatory neurons and inhibitory neurons to the number of all active neurons. In fact, although the E/I ratio was stable (fig.3-j), we observed that in the superficial layers, the neuron density was higher in the occipital to dorsal occipital regions, whereas in the deeper layers, especially layer 6, the neuron density tended to be higher in the frontal side.

**Figure 3.**
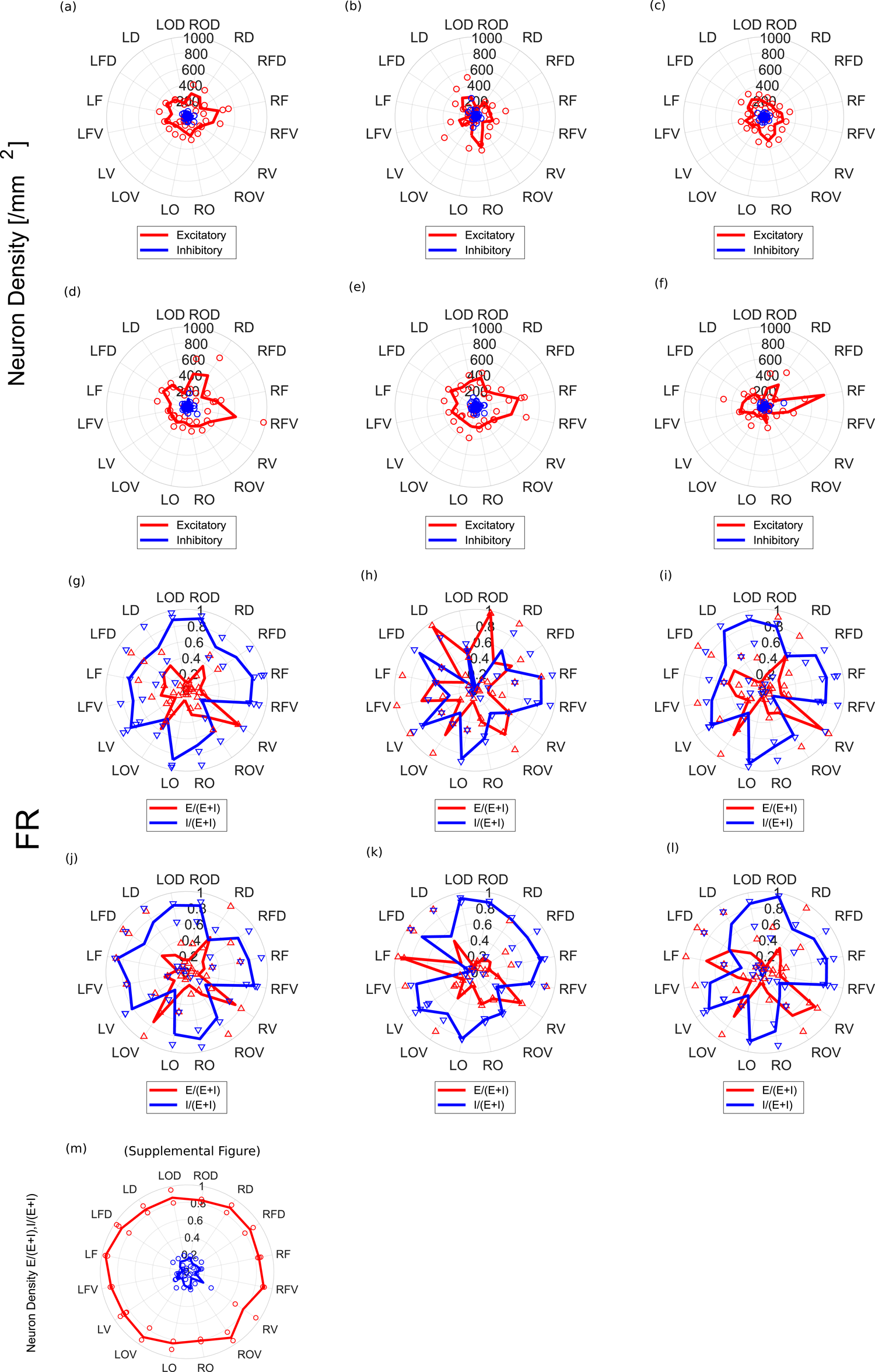
The Most basic physiological metrics, neuron density and firing rates: In all panels, features of excitatory and inhibitory neurons are shown as red and blue lines, and individual data samples correspond to dots. Panel (a) is the radar chart of averaged neuron density included in all layers of the cortical individual brain regions. Panels (b)∼(f) separately express cell densities of individual layers (layer 2, 3, 4, 5, 6). Panel (g) summarizes averaged firing rate in all layers of the cortex, and the firing rates of individual layers (layer 2, 3, 4, 5, 6) are separately shown in panels (h)∼(l). (m) E/I balance calculated as ratio of numbers of active excitatory or inhibitory neurons in the number of all active neurons at individual region groups.

Finally, we observed the firing rate in each brain region. This information beyond structural features is especially important because it is information that cannot be observed from dead slices by staining etc.

Surprisingly, although the firing rate of inhibitory neurons was basically higher than excitatory neurons, a bimodal peak in the firing rate of excitatory neurons was observed around the dorsal (LD,RD) and ventral (LV,RV) regions in both hemispheres. and if we perform statistical test between excitatory and inhibitory neurons, selecting the vicinity of the peaks as target, the firing rate of inhibitory neurons was not significantly greater than that of excitatory neurons (p>0.05, p=0.91, D&V).

Up to this section, we have considered the properties of individual neurons. However, we also need to understand neurons as a network system, in which neurons essentially function by their mutual interactions. Therefore, from the next section, we start to analyze the topological properties as connected networks of nodes, neurons.

### Network metric 1: Degree centrality

In order to move beyond the single neuron level, we will now explore several widely-used network measures. The degree centrality, a basic measure of how central a neuron is among possible paths within a networi, was relatively higher in inhibitory neurons than inexcitatory neurons in most of the regions, in terms both of in-degree and out-degree (in degree:p=0.90, all regions, out degree:p=0.50, all regions).

It has been shown in previous studies that inhibitory neurons are connected to more neurons than excitatory neurons in the somatosensory region [Kajiwara et al., 2021], and a similar trend could be observed at occipital regions (LOD, ROD, LO, RO). Current observation revealed that whether inhibitory or excitatory neurons show higher degree centrality largely depends on the cortical region observed.

### Network metric 2: k-core centrality

In order to check if the tendency observed with degree centrality can be shown with a different centrality metric, we also observed k-core centrality. The radar chart of the k-core of excitatory neuron groups showed that the value at the frontal side (RF, RFV, LF, LFV) is slightly larger than in other regions (corrected p-value=0.16).

### Network metric 3: Controlling ability

Next, we used the feedback vertex set (FVS) index to measure the ability of inhibitory and excitatory cells to control other cells in each local circuit of the brain region. FVS provides a kind of measure of the ability of a neuron to change the intensity of its output while monitoring the surrounding situation by looping the own effects on other cells and returning to oneself.

Here we will briefly explain how to calculate the FVS. In a directed network that can be written with arrows, we focus on a group of nodes. When removal of all incoming connections to the nodes in the group eliminates all directed cycles, the node group is called the feedback vertex set (FVS).

Furthermore, the FVS with the smallest number of nodes is called the minimum feedback vertex set (MFVS). Nodes in MFVS can be considered as driver nodes, and thus we can compare the control ability of the inhibitory neurons and the excitatory neurons by counting the number of MFVS nodes contained in each class of neurons. In our previous study, we evaluated the somatomotor region alone and found that inhibitory cells showed higher controllability than excitatory cells [Kajiwara et al. 2021]. However, MFVS may not be determined uniquely. Therefore, in this study, we employed the weighted minimum FVS (WMFVS), which is the MFVS with the maximum total node weight, where the weight of a node is defined as the sum of edge weights (see below) incoming to the nodes.

In this study, somatomotor regions belong to LFD or RFD. As a result of this cortical wide-area evaluation, it became clear that the dominance of inhibitory to excitatory neurons’ controlling ability shows wide variety among regions [Fig. 4-d]. Here, we could observe relatively higher control ability of inhibitory neurons at LO,LOD,ROD,RO (corrected p-value=0.40).

**Figure 4.**
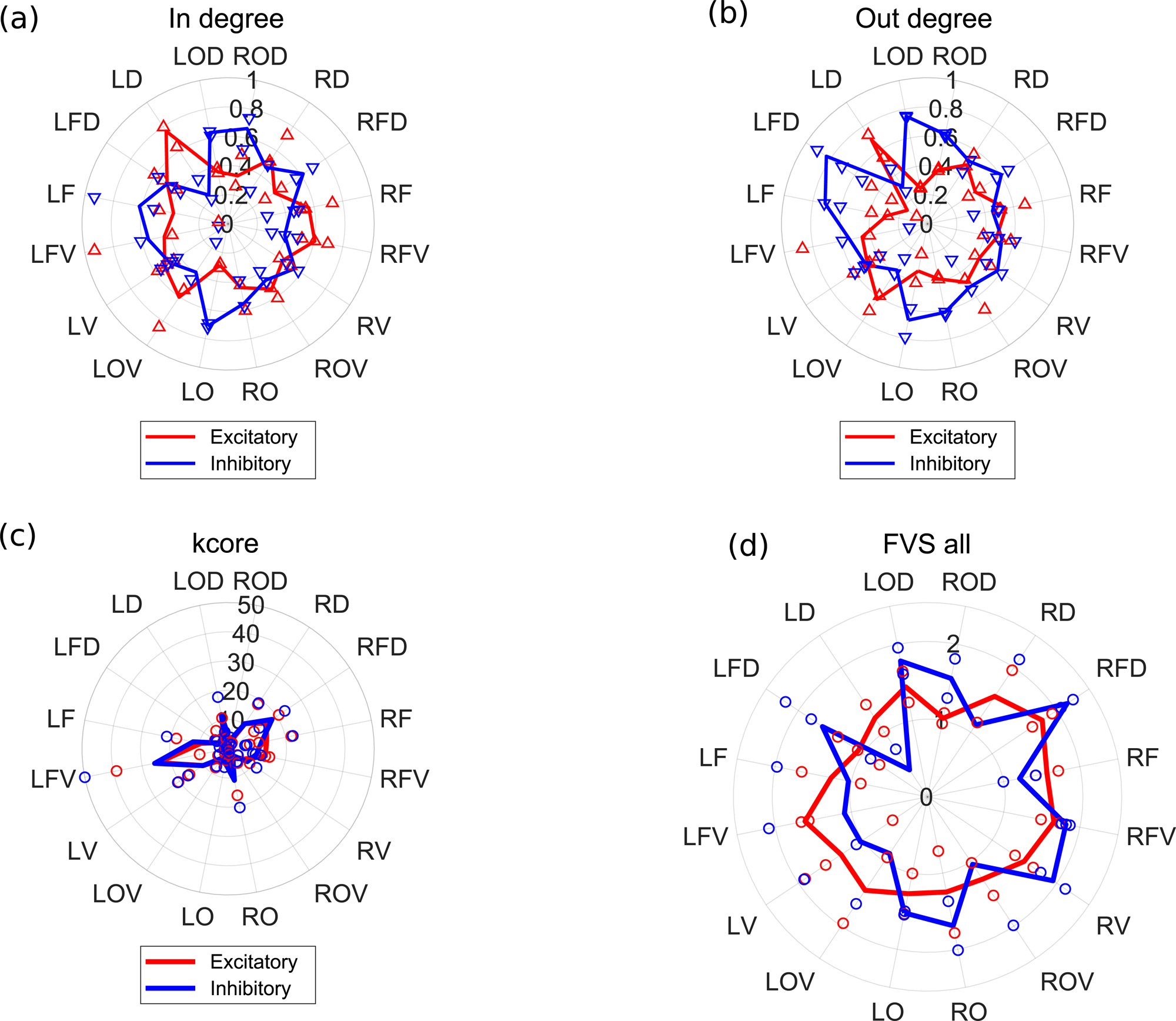
Basic topological properties at individual brain region groups: In panels (a)-(d), red and blue lines show features of excitatory and inhibitory neurons respectively. (a) expresses the number of inward connections, simply called in-degree. (b) also expresses the number of outward connections, called out-degree. (c) is the same radar chart of k-core centralities at individual brain regions. (d) expresses the average FVS (explain here what FVS means) of all layers in the cortex. The meanings of colors of lines and dots are the same as figures 2-4.

### Network metric 4: Weight

Weight is the strength of the connection. All observations up to this point were unweighted (not considering connection strength). And in those observations, the values for inhibitory neurons relatively more often exceeded those for excitatory neurons. We obtained the strength from the amount of information calculated from the spike data using a calculation procedure that shows properties similar to synaptic strength [Song et al. 2005; Nigam et al. 2016].

Next, we decided to observe connection strength (weight). When observing in terms of connection strength (weight), interestingly, the excitatory connections are reliably stronger than inhibitory connections across all cortical regions (corrected p-value=2.5*10^−11^).

Moreover, the connection strength (weight) of excitatory neurons showed a pronounced strength at specific cortical regions, such as LFV, LF, RFV and RF [Fig. 5-a,b] (corrected p-value=0.0011).

**Figure 5.**
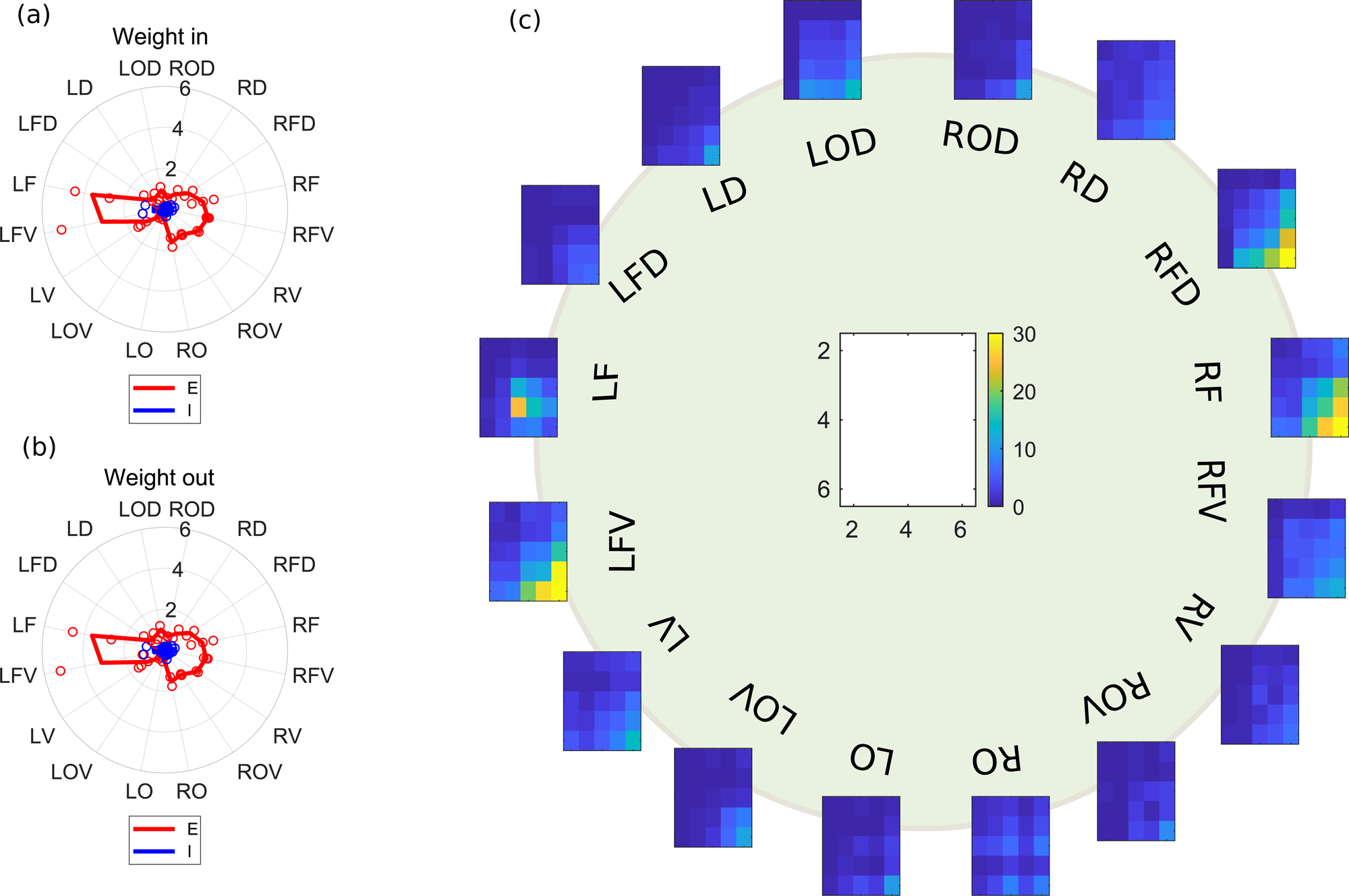
Inter-layer connection strengths: (a) and (b) are similar radar charts of inward and outward connection strengths. **(c)** 16 two-dimensional color maps evaluating the connection strength from input layers (vertical axis) to output layers (horizontal axis) are arranged in a circle. 16 maps correspond to the angles in table1 and many panels in figure 2∼5. When strong connections between input and output layers are strong (weak), it is colored yellow (blue).

These regions are basically the frontal cortex. Recall that the activity quantified with firing rate was strongest in the motor cortex (or somatosensory cortex), which was slightly more dorsal (or occipital) than these areas.

Because current experimentally observed targets are segmented cortical slices, the projecting targets of individual neurons must exist within the observed region. This experimental setting means that the connection strength is defined as mainly and essentially the strength of connections within individual cortical regions. Therefore, simply speaking, we expect that the strong connection within a brain region leads to a strong firing rate at the same brain regions. However, the real results seem to hold a slightly discrepant trend. We will discuss later whether this small discrepancy has any benefit in the brain’s ability to process information effectively over a wide area.

Next, in order to observe the connections between different layers in more detail, we shifted to the observation as a weighted matrix where the horizontal and vertical axes are the indices of the layers.

### Network metric 5: Inter-layer weighted networks

We then observed the connection strengths (connection weights) observed in metric 4, grouped by layer. That is, the connection strength from each of layers 2-6 to each of layers 2-6 was represented as a single matrix for each region. In the individual color maps, the x-axis is for layers where the neurons sending out connections belong, and the y-axis is for layers where the neurons receiving connections belong. As a total, we were able to draw similar connection matrices from 16 regions for the left and right hemispheres [Fig. 5-c], and the angles of 16 maps correspond to the angle used in other figures shown so far.

Although this figure contains too much information to discern entirely, a key feature can be seen: that both of output connections from 6th layers (corrected p-value = 2.4*10^−4^) and input connections to 6th layers (corrected p-value = 0.040) are stronger than the other layers in the frontal and frontal ventral regions which are located around the left and right edges of the figure (yellow regions in fig.5-c). More broadly speaking, we can say that there is a significant tendency for connections with the deep layers to be significantly strong around frontal groups.

## 2. Discussion

In the Results section, we have observed differences among several cortical regions for each single metric. Now we consider the same results from a different perspective. Specifically, the results were reinterpreted by grouping the various metrics observed from the perspective of individual brain regions.

### Main findings

The most interesting properties were observed in the frontal group, where cortical thickness was greater, cell density was higher in the deeper layers, and connection strength tended to be stronger in the deeper layers than in the other brain regions. The cortical region named as the frontal group in this report included the frontal region and frontal side of the secondary motor region.

It has been reported in the past that cortical thickness is greater in this frontal region than in other regions [Fulcher et al., 2019]. We found, in addition, that the trend is observed in the superficial layer thicknesses but not in the deep layer ones.

It was also known in the past that the frontal region is one of the regions with the highest neuron density next to the somatosensory cortex [Keller et al., 2018; Herculano-Houzel et al. 2013]. We also observed neuron density separately by layers. As a result, the layer with significantly highest density in the frontal region was the deeper layer. In the counting of the number of cells detected by staining, the neuron density was greatest in the somatosensory cortex. However, in our result, the neuron density in the frontal cortex exceeded that in the somatosensory cortex because we counted the sufficiently active neurons detected by our electrical activity measurement. In fact, the density of active neurons will not correspond with simple neuron density.

Most interestingly, the connection strength was significantly higher around the frontal group even though the number of functional connections between neurons was not significantly higher in the frontal group [fig4-d]. Connection strength is regarded to be strongly related to response properties to external stimuli [Cossell et al., 2015]. It is said that we can classify connection patterns, connecting to deep layers of the frontal cortex, into multiple modes based on spatial patterns and connecting targets [Gao et al., 2022]. Among the patterns, the wiring extending toward the medial side is especially regarded as a mode of connection pattern that supports the default mode network. In addition, the strong functional connections with the deep layer that we found, in particular, would function to strongly spread inputs, brought in via the spatial global wiring pattern, from/to the deep layers within the frontal area. In the near future, the simultaneous recording of local electrical flow in cellular resolution in frontal region and of global electrical activity spreading along to the global wiring pattern will prove this suggestion more strongly.

The frontal region is also known to be a hub in the extensive network of cortical regions [van den Heuvel, Sporns 2013; Liska et al., 2015], and also intensely interacts with subcortical regions[Oh et al., 2014]. We can offer a possible interpretation of our findings in the context of the current literature. We suggest that the high density of highly active neurons and the strong connection strength in the depths of the network are maintained to serve as an edge (node) in the global network. In particular, we can expect the internal architecture plays an essential role in driving the default mode network observed in the rodent brain [Stafford et al., 2014; Lu et al., 2012; Coletta et al., 2020].

Recently, the relationship between the broad patterns of the default mode network and the cell-by-cell connections in each region have also been discussed [Whitesell et al., 2021]. Our findings add scope to this relationship, given from dynamic spikes of living neurons, to elucidate the facts relating with these many other important findings.

### E/I balance

Next, interesting trends were observed in the regions labeled dorsal region and ventral region. These regions are more active with a relatively higher firing rate of excitatory cells than other regions.

These trends raise an intriguing hypothesis that these regions may be the source of energy that generates high-firing neural activity in the mouse cortex. The slices labeled dorsal in this study are slightly closer to the occipital part of the top of dorsal side, and are located about halfway between the somatosensory and visual areas. It is known that when moving closer to the visual area from this position, the position becomes a slice that retains connections with subcortical thalamus [MacLean et al., 2006]. It is important to note that in our results, the firing rate of excitatory neurons does not increase as one approaches the occipital visual area from the dorsal region. In other words, the higher firing seen in our dorsal area does not reflect the effect of the interaction loop with the thalamus, but rather the property of the internal local circuits.

By the way, even if the E/I balance differs from region to region, we can say that the E/I balance is well maintained in cortical regions as a whole. There is a differential trade-off between speed and specificity in increasing the proportion of excitatory and inhibitory neurons [Sadeh et al., 2015]. In this situation, optimal balance of excitatory and inhibitory synapses may allow an asynchronous irregular network to sustain a homeostatic state, which could be re-activated by external stimuli [Vogel et al., 2011]. Such an arrangement could enables the system to generate sharp responses, to generate oscillations, and also to achieve optimal control gain and dynamic range [Isaacson, Scanziani, 2011]. We used relatively young mice, 3weeks old. This age range could reflect their importance in ongoing pattern formation in the process of developing neural circuits [Sadeh, Clopath, 2021]. The experimental results in our study are consistent with the fact that, if we observe activities of living neurons as the whole of cortical regions, the contributions of excitatory and inhibitory neurons are highly balanced.

### Future works

As described above, the present study has derived from the data some very interesting trends related to brain nonuniformity, which compensate for information that is not available from information on regional connections and structural cellular distribution previously provided from different recording modalities. However, since the scope of the analysis was limited within the cortex, it wil beimportant in the future to develop the analysis to include subcortical regions as well. In fact, it is known that there are subcortical areas where cell densities are higher than in the cortex [Keller et al., 2018]. More analysis of the same cortical data after parcellating with a cutting edge atlas will also be important in the future to compare with results at more specific brain regions. It is also interesting to contrast the microconnectome obtained by our method with the structural microconnectome, which is obtained using SEM etc [Motta et al., 2019]. It is also important to compare the results with the non-uniformity of genes expressed in the various brain regions [Waterston et al., 2002; Lein et al, 2007; Fulcher et al, 2019]. This study took three years to accumulate more than two data points from 3∼5 week old mice, allowing us to check for repeatability in each area. Although significance was observed in the trends described above, it will be important to increase the number of samples and to obtain data from elder mice in the future. In any case, there has been no study that systematically measures activity at ms temporal resolution from the whole cortex and discusses its activity and interaction network as in this study. Therefore, we expect the data and findings obtained in this study should be extremely valuable. This study may also be useful for improving future mathematical models of the brain and for advancing the systematic understanding of disease through animal models.

## Materials and Methods

### Physiological experiments

The present study acquired acute brain slices from 16 groups of cortical regions of the cerebral cortex and simultaneously measured electrical activity from a large number of neurons within the brain slices [figure 2-a; table1]. The series of experimental protocols, including 3D scanning and immunostaining before and after that electrical activity measurement, followed the experimental protocols used in previous studies that have already been published studies including a video journal explaining it in detail [Ide et al. 2019; Kajiwara et al. 2021; Shirakami et al. al., 2022]. All animal experiments were performed according to animal experiment protocols approved by the Kyoto University Animal Committee. All mice used were female C57BL/6J mice (n=37, 3-5 weeks old).

The thickness of the cortical slices was 300 μm, and we used a Multi-elecord array (MEA) system (Maxwell Biosystem, MaxOne) for simultaneous measurement of electrical activity. During the measurements, we refluxed an artificial cerebrospinal fluid solution saturated with 95% O2/5% CO2.

Also, we performed spike sorting (Spyking Circus software) on the time series obtained from the electrical measurements to identify ∼1000 neurons; the short distance spacing between the electrodes of the MEA system (15 µm) allows us to estimate the spatial position of the neurons very accurately.

Before and after the electrical measurements, we measured 3D scans of the brain surface [Ide et al., 2019], and by comparing and superimposing the scan data with the MR images recorded before the brain was extracted, we were able to accurately define the slice positions of individual brain regions with less than one slice resolution (Figure 1a).

The cerebral cortex has a “baumkuchen”-like pattern of six overlaid sheets in the depth direction. Physiologically, each of these sheets are called layers. The layers are named layers 1-6 in order from the surface of the brain toward its depth. The layer architecture could be visually observed by immunostaining using NeuN and GAD. In this study, layers 2-6 are discussed because layer 1 is not included in the measuring area of some samples in the electrical measurement.

### Defining connectivity reflecting causal neuronal interactions

This study used transfer entropy (TE) to quantify the causal interactions between neurons (Figure 1d). TE has been recognized by various research groups as an excellent and standard metric to assess the causal interactions of neural activity and has historically accumulated a large number of studies [Wibral et al. 2013; Lizier et al. 2008; Garofalo et al. 2009; Stetter et al. al.2012; Orlandi et al. 2013].

Networks of causal interactions are broadly referred to as effective networks [Friston 1994; Aertson, 1989]. We also prepared a new network inference method enabling us to categorize excitatory and in-hibitory connections, and neurons. The details of the analysis methodology used in this study are summarized in Kajiwara et al. 2021. The source code is also available on git-hub (https://github.com/Motoki878).

## Network metrics

### Degree

Degree is a metric defined for each node and is the number of edges connecting to each node. In a directed graph with oriented edges, there are incoming edges and outgoing edges, so two indices, in-degree and out-degree, are defined for each type vertex (Figure 4-a). The in-degree is the number of edges coming into the vertex, and the out-degree is the number of edges going out from the vertex. The distribution of frequencies of vertices with similar degrees is called degree distribution, and is frequently used to characterize structure of networks.

### Weight

Weight refers to the strength of the connection, and here it is calculated by normalizing the amount of information calculated from the real spike data by the amount of information obtained when the time sequence of the spikes is shuffled. In other words, weight assesses the rate at which firing between two neurons occurs causally within a few milliseconds.

Past studies have reported that this metric shows statistically similar trends to synaptic connections [Song et al. 2005; Nigam et al. 2016; Kajiwara et al. 2021]. It should be noted that if one neuron outputs more neurons, the amount of information sent to the individual destination neurons is smaller.

### k-Core

Next, we used the k-Core index as another representative measure of centrality along with degree [Seidman, 1983]. k-Core is determined by removing a node with 0 connections (isolated vertex) to a node with 1 connection, to a node with 2 connections, and so on from the original connection structure. In general, k-Core is defined as a partial network consisting only of nodes that (1) remain after removing nodes of order less than k, and (2) have a number of connections greater than k. To define k-Core in consideration of both rules (1) and (2), it should be noted that k-Core does not simply correspond to a network consisting of nodes with degree greater than or equal to k. many cases.

### FVS

A set of nodes in a directed graph (or network) is called a feedback vertex set (FVS) if removal of the incoming edges to the set eliminates all directed cycles. It is known that for reasonably wide classes of linear and non-linear systems, statically and periodically steady-states of the whole network are determined by specifying states of the nodes in an FVS [Akutsu, et al., 1998; Mochizuki et al., 2013].

The minimum feedback vertex set (MFVS) is the FVS with the minimum number of nodes and can be regarded as a minimum set of driver nodes. However, MFVS may not be uniquely determined in many cases. In order to cope with this issue, the weighted minimum FVS (WMFVS) was introduced by making use of weights assigned to nodes, where WMFVS is the MFVS with the maximum total node weight [Li et al., 2021]. It is expected that WMFVS is uniquely determined if various weights are assigned to the nodes. Although computation of both MFVS and WMFVS is theoretically difficult (NP-hard), these can be computed efficiently for moderate size networks by using integer linear programming [Li et al., 2021].

## Acknowledgments

MS is supported by several MEXT Grant-in-Aid for Scientific Research (B) (20H04257) and the Leading Initiative for Excellent Young Researchers (LEADER) program. The MRI experiments of this work were performed in the Division for Small Animal MRI, Medical Research Support Center, Graduate School of Medicine, Kyoto University, Japan. We acknowledge John M. Beggs, Zakian Doris fruitful comments on this manuscript, and all support from Innovative Support Alliance for Life Science, and the Hakubi Center in Kyoto university to complete this study.

